# Hyperporous encapsulation of microbes for whole cell biocatalysis and biomanufacturing

**DOI:** 10.1101/2024.06.18.599529

**Authors:** Jingyi Zhang, Keziah Chang, Joyce Tay, Elaine Tiong, Elena Heng, Theresa Seah, Yi Wee Lim, Guangrong Peh, Yee Hwee Lim, Fong Tian Wong, Cyrus W. Beh

**Affiliations:** Bioprocessing Technology Institute, Agency for Science, Technology and Research (A*STAR), 20 Biopolis Way, #06-01 Centros, Singapore, 138668, Singapore; Institute of Molecular and Cell Biology (IMCB), Agency for Science, Technology and Research (A*STAR), 61 Biopolis Drive, Proteos #07-06, Singapore, 138673, Republic of Singapore; Department of Biomedical Engineering, National University of Singapore, 4 Engineering Drive 3, #04-08, Singapore 117583, Singapore; Institute of Sustainability for Chemicals, Energy and Environment (ISCE^2^), Agency for Science, Technology and Research (A*STAR), 8 Biomedical Grove, Neuros, #07-01, Singapore, 138665, Republic of Singapore

## Abstract

Compared to traditional synthetic chemical processes, biocatalysts offer a more sustainable and eco-friendly approach to producing complex molecules. In particular, whole-cell biocatalysts boast numerous advantages, including scalable, self-containing co-factor recycling systems, the use of cost-effective raw materials, and reduced purification costs. However, challenges arise when working with microbial consortia for biotransformation cascades. Our encapsulation strategy addresses these challenges by controlling microbial cell populations through physical constraints, offering a promising approach in biomanufacturing.

In this work, we describe the immobilization of cells in a hyper-porous hydrogel block, which provides ample nutrient access while simplifying media changes. We encapsulated *E. coli* cells in a hydrogel matrix with suitable mechanical properties, effectively limiting their proliferation while sustaining recombinant GFP production. Furthermore, we successfully maintained different microbial strains spatially in a single porous hydrogel block for at least 10 days, demonstrating the potential of this method for achieving stable co-culture. Finally, we demonstrated the application of immobilized *E. coli* for co-culture fermentation. The immobilization of *E. coli* heterologously expressing RadH halogenase significantly improved the efficiency of genistein halogenation in a co-culture with genistein-producing *Streptomyces* compared to its non-immobilized counterpart.

## Introduction

Compared with traditional chemical processes, biocatalysts offer opportunities for sustainable and environmentally friendly manufacturing of complex molecules. With its high specificity and mild reaction conditions, there is less chemicals, waste and energy required.^1, 2^ While industrial practice has tended to favor immobilized enzymes over whole cell biocatalysts in recent years,^2^ the latter still offers some advantages over cell-free enzymes.^3^ For example, cell-free systems often require use of purified enzymes, which increases the time and cost of production.^3^ In many reactions, it is also necessary to supply expensive co-factors, or otherwise develop recycling systems, further increasing production costs.^2^ On the other hand, whole-cell biocatalysts contain inherent co-factor recycling systems, simplifying the process. The cells can also convert cheap raw materials into high-value chemicals that have applications in pharmaceutical industry.^3-5^ For these reasons, research into whole-cell biocatalysts have remained active.^4-6^

These advantages notwithstanding, several challenges persist for the adoption of whole-cell biocatalyst. Traditionally, a single industrial strain is engineered to incorporate the entire synthetic pathway,^7^ though the heavy gene expression burden can result in reduced efficiency and robustness.^8^ In nature, microbial consortia comprising different species accomplish the task of performing difficult chemical tasks by division of labour. Inspired by this, some groups have designed co-culture systems with a mixture of prokaryotic and eukaryotic organisms.^6, 7^ However, a stable co-culture is difficult to achieve, and may involve elaborate strain engineering to create a mutualistic relationship between the consortia members.^7^ While possible for a two-organism system, such engineering can clearly become intractable when the types of organisms increase.

In this report, we describe our work developing an encapsulation strategy to achieve control of microbial cell populations by physical means. Encapsulation of *Escherichia coli* (*E. coli*) cells in a hydrogel matrix with suitable mechanical properties constrains their proliferation, reaching a plateau after a few days. Our method results in the immobilization of the cells in a hyper-porous hydrogel block, which permits ample access to nutrients by the cells, while enabling media changes to be easily accomplished. We also showed that different microbial strains can be maintained in a single porous hydrogel block for at least 10 days, thereby demonstrating the potential of this method for achieving stable co-culture. Finally, we demonstrated that these immobilized *E*.*coli* retained their capabilities for recombinant protein expression and as whole cell biocatalysts.

## 2. Materials and Methods

### 2.1 Materials

Gelatin from porcine skin (gel strength 300, type A) was purchased from Sigma Aldrich. Microbial transglutaminase (mTG) was purchased from Modernist Kitchen. Phosphate buffered saline (PBS) was purchased from Gibco. Collagenase from Clostridium histolyticum type IV and lithium phenyl (2,4,6-trimethylbenzoyl) phosphinate (LAP) were purchased from Merck. Hanks’ Balanced Salt Solution (1X) (HBSS) and Nunc Lab-Tek chamber were purchased from Thermo Scientific. Mueller Hinton II Broth (MHB) was purchased from BD Biosciences. Organic solvents (DMSO, ethanol, acetone, acetonitrile) were purchased from Sigma Aldrich. 96-well clear flat bottom polystyrene TC-treated microplates were purchased from Corning. Petri dish (94 mm x 16 mm), not-treated, polystyrene was obtained from Greiner. Gelatin methacryloyl (gelMA) (degree of methacrylation: 75 – 85%) was purchased from Gelomics. Samples were imaged using Carl Zeiss LSM 700 confocal microscopy. Particles were prepared using handheld extruder equipped with a coarse 3mm holes and mesh with pore size (diameter) of 340 µm. Biowave Co8000 cell density meter was used to measure the OD_600_ readings.

### 2.2 mTG toxicity test with *E. coli* in Mueller Hinton Broth (MHB)

MHB buffer (1.76% w/v) was prepared by dissolving MHB powder in MilliQ water. Using a stock solution of mTG in MHB buffer (10% m/v), eight different concentrations (0.07813%, 0.156%, 0.3125%, 0.625%, 1.25%, 2.5%, 5%, 10% m/v) were prepared via serial dilution. *E. coli* containing a plasmid for expression of enhanced green fluorescent protein (eGFP) was grown in Luria-Bertani (LB) with resistance at 37°C and 200 rpm until it reached OD_600_ of 0.7. The E. coli stock was then diluted a further 10, 000x in each concentration of mTG solution in addition to a control group (pure MHB) with antibiotics included. In a 96-well clear bottom microplate, 100 µL of this solution was added to each well and the baseline OD_600_ reading measured. The plate was then incubated at 37°C and 100 rpm for 18 h before a final OD_600_ reading was taken.

### 2.3 Preparation of Hyper-porous Hydrogel

20% w/v gelatin solution with 2.5 % w/v mTG was prepared at 37°C and mixed thoroughly to achieve a homogeneous solution. After cooling, hydrogel was passed through the extruder multiple times to form microparticles. The microparticles were then rinsed with mTG solution before submerging in the mTG solution overnight, resulting in the formation of a hyperporous, granular block scaffolds.

Granular blocks were washed in MilliQ water and shaken in individual flasks at 37°C and 100 rpm. Each block was cultured in 20 mL LB with 5, 10 or 20% v/v organic solvent (DMSO, ethanol, acetone, acetonitrile) containing antibiotics and 0.1 mM IPTG. Controls of blocks cultured in 20 mL LB with antibiotics and 0.1 mM IPTG were conducted as well. Dimensions of the blocks (length, width and height) were taken daily with vernier callipers to calculate the volume.

### 2.4 Encapsulation of *E. Coli*

For *E. coli* encapsulation, gelatin and mTG solutions were prepared in LB broth containing IPTG and antibiotics. At 37°C and 220 rpm, eGFP *E. coli* was seeded in LB with antibiotics and sub-cultured the next day in 5 mL LB with antibiotics until it reached OD_600_ of 0.4. Cells

were then induced with 0.1 mM IPTG at 37°C and 220 rpm for 1 h. The porous granular blocks were then prepared as described in Section 2.3.

### 2.5 Release of Cells from Encapsulating Hydrogel

Granular blocks were each rinsed with 100 mL MilliQ water to remove free bacteria. To release *E. coli* cells from the hydrogel, each granular block was shaken with 1 mL 0.5 mg/mL collagenase solution at 37°C and 1500 rpm for 45 mins. The collagenase solution was prepared by dissolving powdered collagenase in Hanks’ Balanced Salt Solution (HBSS). The subsequent cell solution was neutralized with 2 mL LB and spun down at 3000 rpm for 10 mins. The supernatant was discarded, and the cell pellet was resuspended in 5 mL LB. For colony-forming units (CFU) counts, cells were serially diluted to concentrations ranging from 10^−6^ to 10^−11^, plated on LB agar plates with antibiotics and incubated at 37°C overnight. Colony counts were taken the next day and results reported as logCFU values for each individual block.

### 2.6 Hydrogel stiffness experiments

10% w/v gelatin (with 2.5% w/v mTG) and 20% w/v gelatin (with 2.5% and 5% w/v mTG) solution were prepared by dissolving gelatin and mTG in PBS. Each resulting solution was allowed to crosslink at 37°C for 1 h. After crosslinking, the gel was left to equilibrate to room temperature for an additional hour. A biopsy punch was used to create three samples per condition. Subsequently, compression tests were performed on these discs with varying crosslinking conditions (Instron 3340 frame). The stiffness (Young’s modulus) was quantitatively measured by calculating the slope of the linear portion of the stress-strain curve obtained during the test.

### 2.7 Monitoring Growth of Encapsulated Cells

Hydrogels of different stiffness containing eGFP-expressing *E. coli* are prepared as described, and cultured in LB media, 100 µg/mL ampicillin, and 0.1 mM IPTG. At predetermined time points, the hydrogels were retrieved from the culture flasks, and rinsed with deionized water. Fluorescence and brightfield images were used to monitor the diameter of the colonies over time. Each condition contains 3 samples, with 20 colonies sized for each sample. Long term culture was performed using 20% gelatin crosslinked with 2.5% mTG. The samples were cultured as described in the previous paragraph, and colony size was monitored for 51 days.

### 2.8 Co-Culture of Microbial Strains

Co-culture of eGFP- and DsRed-expressing *E. coli* strains was performed by suspending the microbes in LB media containing 20% gelMA, 2 mM LAP, and 0.1 mM IPTG, and forming porous hydrogel blocks as previously described.^9^ The samples were then imaged at various time points on the confocal microscope. The particle boundaries were determined using a combination of brightfield images and scattering by the hydrogel matrix.

### 2.9 Halogenation by Encapsulated Microbes

*Streptomyces* sp. A1301 was seeded for 3 days in SV2 media, followed by sub-culturing in 50mL CA10LB^10^ fermentation media. 3mL culture was collected at 24 h for LCMS analysis before the addition of immobilized *E*.*coli* expressing RadH.^11^ *E*.*coli* was seeded in LB with resistance and sub-cultured the next day in 20 mL LB with resistance until it reached OD_600_ of 0.4. Cells were induced with 0.1 mM IPTG and encapsulated before adding to the 24 h fermentation culture of *Streptomyces* sp. A1301. All samples were done in triplicates. Samples were taken at day 7 for analyses.

## 3. Results

### 3.1 Encapsulation Protocol

Encapsulation begins by dispersing microbial cells in warm gelatin. After the gelatin cools to room temperature, the hydrogel is passed through a filter mesh to form hydrogel microparticles. The microparticles were then washed with a microbial transglutaminase (mTG) solution at the stated concentration, and cast in a chamber at room temperature overnight. Enzymatic crosslinking takes place in the particles and at the particle-particle interface, resulting in a covalently crosslinked hyper-porous hydrogel block that remains intact even at the *E. coli* incubation temperature of 37 °C, which is higher than the melting temperature of gelatin. The porosity allows media exchange to occur efficiently simply by immersion in fresh media.

The stability of the hydrogel construct is an important consideration in encapsulation. In particular, for biomanufacturing of small molecules, stability in organic solvents (e.g. DMSO, ethanol, acetone, acetonitrile)^12^ would enable biocatalytic processes at much higher substrate concentrations, achieving desired process intensification, and also promote longevity of the biocatalysts. Our preliminary results indicate that the hydrogel blocks are stable in up to 20% organic solvents with no changes apart from ∼5% shrinkage after 6 days incubation.

### 3.2 Cell Viability After Crosslinking

Crosslinking of the hydrogel involves mTG, which can affect the viability of the *E. coli* due to its protein crosslinking ability.^13-16^ To assess the biocompatibility of our process, we measured cell density (OD_600_) after 18 hours of incubation under various mTG conditions. At the typical concentration of 2.5% we use for our preparations, there was no significant reduction in OD_600_.

To assess viability, we measured the CFUs of the cells before and after encapsulation. Cells were released by digesting the gelatin with collagenase. With an input of 5 logCFU/mL, we are able to achieve > 9 logCFU/mL after overnight encapsulation, proving that the cells are still viable.

### 3.3 Achieving Stable Encapsulated Cell Population

Our approach uses physical means to constrain the growth of cells. For this reason, we hypothesized that the material stiffness will affect the cell proliferation. To verify this, hydrogels containing *E. coli* with different stiffness were prepared by varying either/both gelatin and mTG concentrations. Samples were thus imaged on a confocal microscope at fixed intervals to measure the colony sizes (Figure 3A). Samples were also mechanically tested by compression to determine the Young’s Modulus.

**Figure 1.**
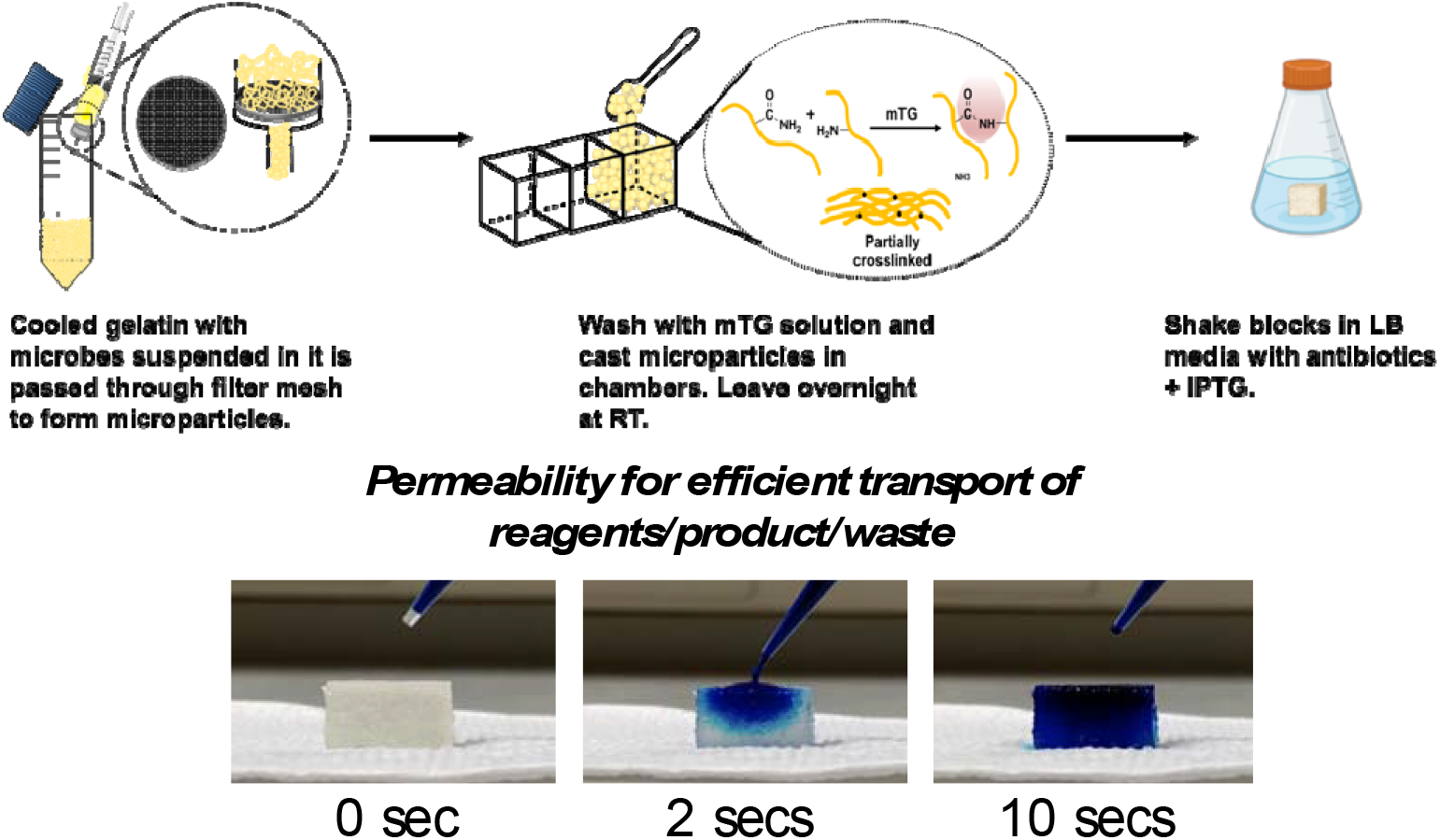
Preparation of porous hydrogels from microbe-laden hydrogel microparticles.

**Figure 2.**
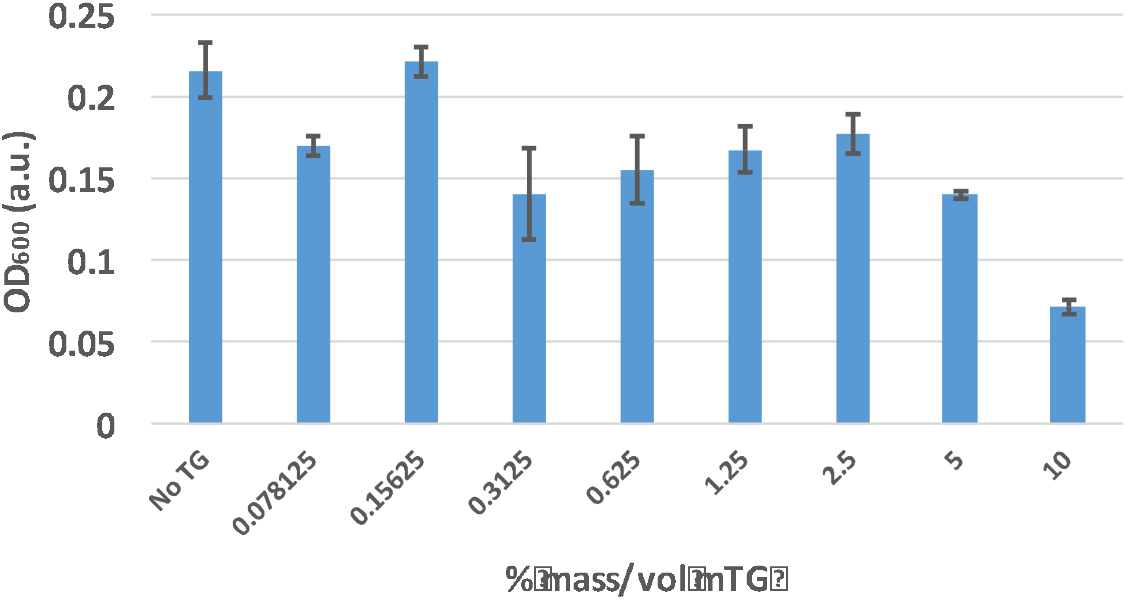
OD_600_ measurement of *E. coli* incubated with different mTG concentrations in MHB media. N = 6 per concentration.

**Figure 3.**
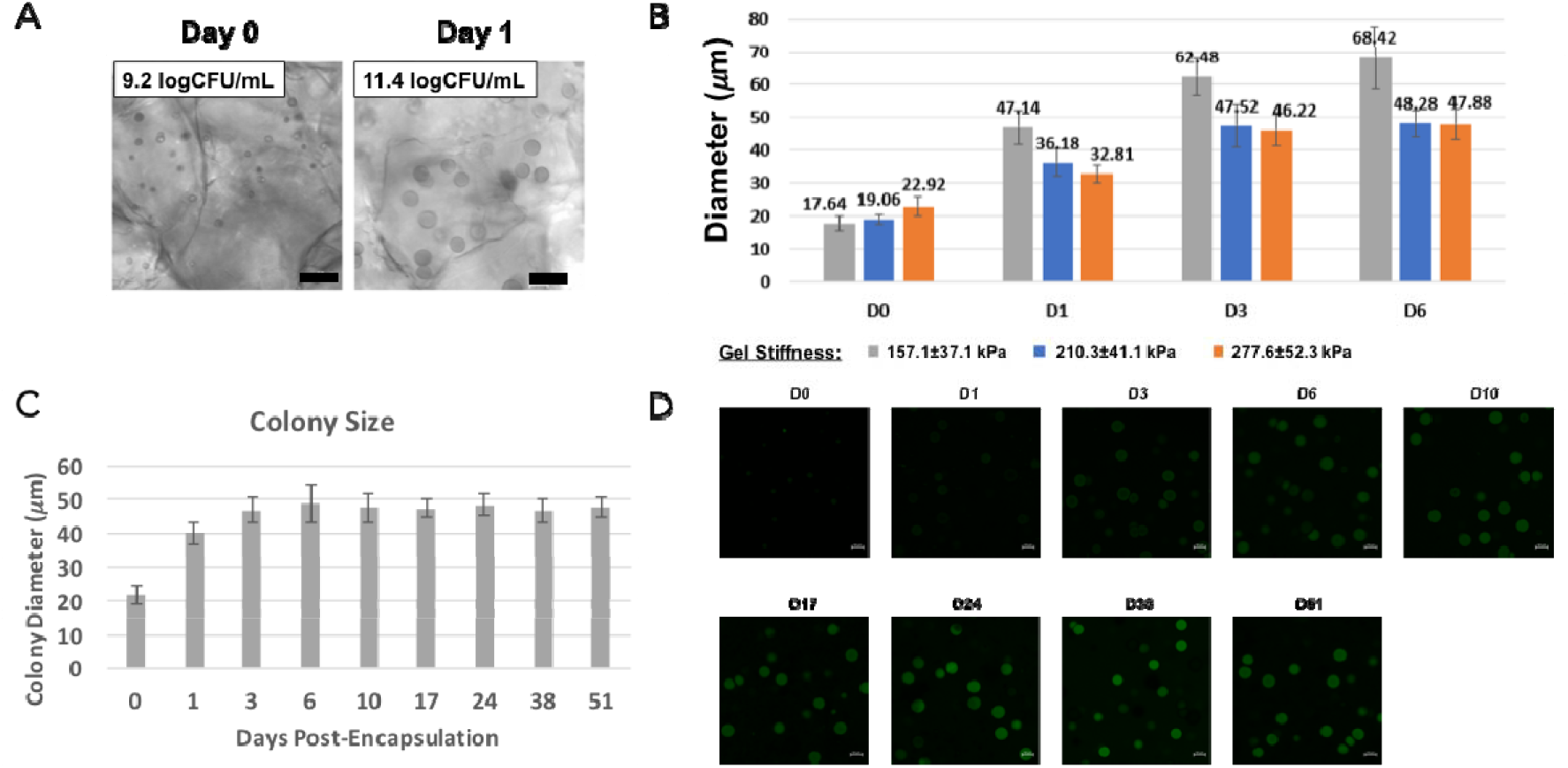
Encapsulation of *E. coli* in hydrogel. (A) The cells form visible colonies, which correlates with the cell number. Scale bar = 100 µm (B) Hydrogels of different stiffness were prepared with *E. coli* encapsulated within. Colonies of cells in the 157 kPa gel (gray bars) increased throughout the experiment, and the hydrogel degraded on Day 7. On the other hand, the colony sizes in the stiffer gels stabilize around Day 3. D0 corresponds to the sample after overnight crosslinking. (C) Using the 210 kPa gels, we were able to achieve prolonged culture of cells, with colony sizes staying largely unchanged between days 6 and 51. (D) We also observed almost no eGFP expression during the fastest colony growth from D0 to D3. eGFP expression gradually increased after D3 and continued up to D51.

We found that the sample with the lowest stiffness indeed resulted in the fastest cell growth (Figure 3B). Furthermore, despite the relatively close gel stiffness, the 157 kPa gel degraded 7 days post-crosslinking. The size of colonies also increased continuously, which may have contributed to the degradation of the hydrogel by rupturing the microparticles. On the other hand, the colonies in the stiffer gels grew more slowly, and reached a plateau around Day 3. This may have prevented the disintegration of these gels.

Using the 210 kPa gels, we encapsulated *E. coli* and cultured the hydrogel block on a shaker with daily LB media change over a prolonged period. The samples were retrieved for imaging at fixed intervals (n = 3, 20 colonies sized for each sample at each time point). We found that the cell growth was highest immediately post-encapsulation, but stabilized from around Day 6 (Figure 3C). eGFP signal, on the other hand, was low initially, and gradually increased from around Day 6 (Figure 3D). At Day 51, eGFP signal can still be observed. This suggests that cells direct resources away from production during the growth phase and is consistent with previous reports.^3, 17, 18^ Observations of consistent production of eGFP, while colony diameter remain constant, suggests that physical constraints of our encapsulation strategy have redirected resources from biomass production to recombinant protein production. This implication could significantly impact biotransformation efficiencies.

### 3.4 Stable Co-Culture of *E. coli*

For multi-step biotransformation cascades, it may be advantageous to divide the synthetic pathway into multiple strains to reduce the workload. However, it is challenging to achieve stable co-culture of multiple strains in a single pot. For example, in our initial experiments with culture fermentation, the ratios between strains were significantly inconsistent, ranging from 1:1 to 20:1 within 24 hours. We performed a co-culture of *E. coli* expressing either eGFP or DsRed by separately encapsulating each strain in particles, then crosslinking a blend of the two particle types into a single hydrogel block. This ensures that the two strains are compartmentalized in their respective particles.

Over a period of 10 days, each strain remained within its own particle, with no evidence of cross-ingression. More than a simple single-pot reaction, this configuration brings the two strains into close proximity, allowing efficient transport of substrates and intermediates between the strains, and can also result in high effective local concentration of various metabolites. This may be of particular interest to processes for which short-lived intermediates have to be quickly processed by downstream biotransformation reactions.

### 3.5 Enzymatic Halogenation Reaction

To demonstrate the importance of immobilisation in a co-culture biotransformation, we also examined halogenation of genistein by RadH halogenase^11^ heterologously expressed in *E*.*coli* in a co-culture fermentation with *Streptomyces*. Here, we compared the levels of halogenation with and without immobilisation (Table). As before, *E. coli* is encapsulated in our hyper-porous hydrogel block, and co-cultured with non-immobilized *Streptomyces*. We were able to observe significant conversion of genistein (**1**) into its chlorinated products (**2** and **3**), as determined by LCMS-QTOF after 7 days of fermentation. These results show that diffusion of small molecule substrates and products through the hydrogel matrix is possible, and that the encapsulated microbes continue to function as whole-cell biocatalysts.

**Table 1.**
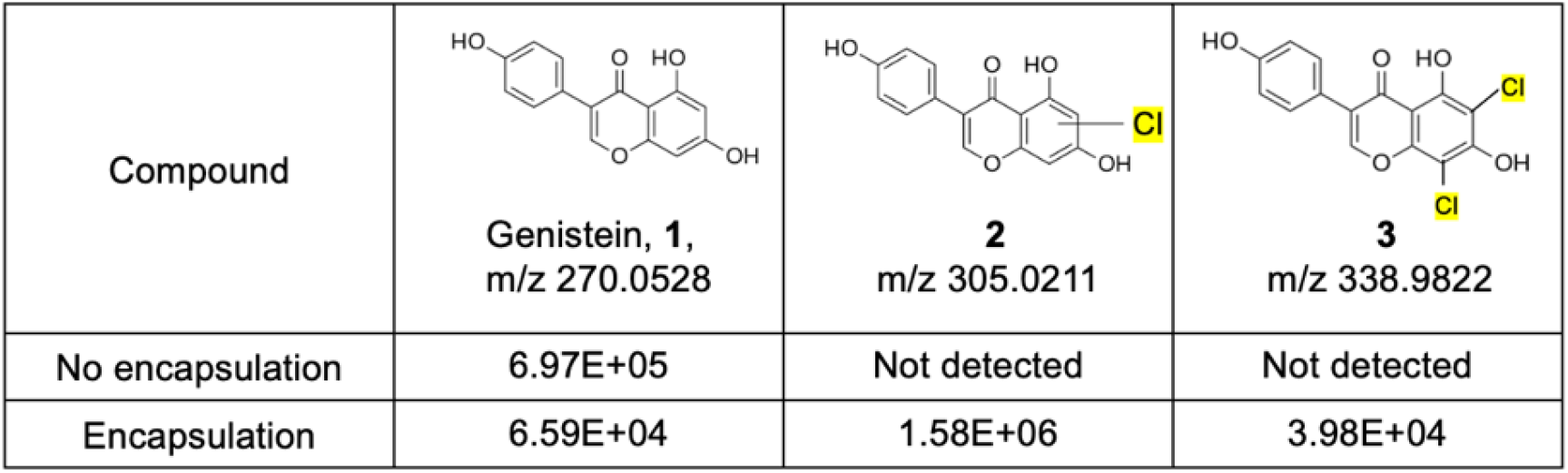
Comparison of chlorination of genistein (**1**) by encapsulated or free *E*.*coli* expressing RadH halogenases in a co-culture fermentation at day 7. (Peak intensities are given from LCMS-QTOF analyses)

| Compound | Genistein, **1**,<br>m/z 270.0528 | **2**<br>m/z 305.0211 | **3**<br>m/z 338.9822 |
| --- | --- | --- | --- |
| No encapsulation | 6.97E+05 | Not detected | Not detected |
| Encapsulation | 6.59E+04 | 1.58E+06 | 3.98E+04 |

While the current experiment uses free *Streptomyces* with encapsulated *E. coli* in shake flasks, we envision that co-culturing of the two microbes at an optimized ratio, in a single porous block (as in Figure 4), will significantly enhance the biotransformation process. Additionally, immobilizing the microbes will allow for continuous biomanufacturing and facilitate the transition from growth to production phases. This setup will also enable us to optimize the biotransformation reaction, including media optimization for production rather than growth (i.e. organic solvent or minimal simple media). This will allow for further intensification of the reaction while simplifying downstream processing.

**Figure 4.**
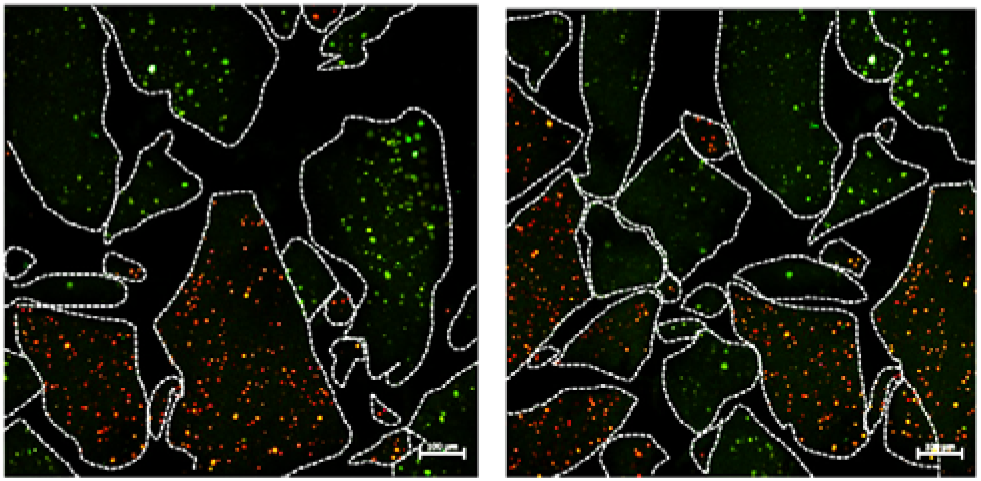
Encapsulated *E. coli* colonies remain compartmentalized to their own particles after at least 10 days of culture. White dashed lines demarcate each particle boundary. Scale bar = 100 µm

## Conclusions

In conclusion, we have demonstrated a facile method for encapsulating fast-growing *E. coli* in a hyper-porous hydrogel block. The porosity allows easy access to nutrients in media, clearance of metabolites, and transport of enzymatic substrates to the cells. The encapsulated cells remain viable, but their proliferation is physically constrained by our hydrogel matrix. By tuning the mechanical properties of the matrix, it is also possible to redirect resources from biomass production and achieve stable culture for at least 50 days. Furthermore, this strategy can be used to facilitate stable co-culture of different strains in a single hydrogel structure, opening up possibilities for efficient biotransformation cascades to be performed. Lastly, we demonstrate that the encapsulated microbes remain enzymatically active, as evidenced by the successful halogenation of the genistein substrate. While we have shown a proof-of-principle in this study, further optimization of the various processes is currently on-going to fully realize the potential of our method in enhancing biocatalytic productivity.

## Acknowledgements

The authors gratefully acknowledge funding support from Agency for Science, Technology and Research (A*STAR) (C2333017003 and C2333017004) and A*STAR Graduate Academy (to K.C. and T.S.).

## Author contributions

C.W.B.: Conceptualization, methodology, software, data curation, formation analysis, visualization, writing - original draft-writing, writing - review and editing

J.Z., K.C., J.T.: Investigation, formal analysis, writing – original draft

T.S.: Investigation, formal analysis

F.T.W.: Conceptualization, investigation, methodology, writing - original draft-writing, writing - review and editing

G.P., E.T., E.H., Y.W.L.: Investigation, methodology, formal analysis

Y.H.L.; Conceptualization, Data analysis, writing - review and editing.

